# NPDS Toolbox: Neural Population (De)Synchronization toolbox for Matlab

**DOI:** 10.1101/2021.07.13.452294

**Authors:** Mohammad Mahdi Moayeri, Mohammad Hemami, Jamal Amani Rad, Kourosh Parand

## Abstract

The study of synchronous or asynchronous in (stochastic) neuronal populations is an important concept both in theory and in practice in neuroscience. The NPDS toolbox provides an interactive simulation platform for exploring such processes in Matlab looking through the lens of nonlinear dynamical systems. NPDS includes two main components: neural population (de)synchronization, and neural dynamics. One can investigate distribution controls on various neural models such as HH, FHN, RH, and Thalamic. Also, it supports many numerical approaches for simulation: finite-difference, pseudo-spectral, radial basis function, and Fourier methods. In addition, this toolbox can be used for population phase shifting and clustering.

## 1. Introduction

Synchronization is a vital process in most complex dynamic systems, especially in neural networks within the brain. In fact, various cognitive functions such as decision making, learning, perception, memory, etc. are the result of the synchronization of neuronal populations [1, 2, 3, 4]. However, abnormal and excessive neuronal synchrony in different parts can be one of the reasons for some neurological disorders such as Parkinson’s disease or epilepsy [5]; thus, a fine balance between synchronization and desynchronization is functionally and behaviorally important [6]. Using the control mechanisms in the neural population is a practical approach to modulate seizure activity which is considered by the researchers. Among various types of control strategies [7, 8, 9], neurons phase distribution control [4, 10] is one of the suitable options because it has remarkable features; for instance, in this method, this control takes the analysis of high-dimensional systems more tractable, which reduces the time of solving the problems and improves the performance of the control. It also optimizes the energy consumed by defining a proportional control. Another feature of this method is its applicability to any experimental and neuronal model with respect to its phase response curve (PRC), which makes this technique applicable to any system having a (de)synchronization challenge [10, 4]. However, according to Moehlis et al. [4, 10] and our opinion, the performance of control strategies, especially this phase distribution control, relies heavily on numerical methods to simulate these dynamic nonlinear models, so that using an advanced and more accurate numerical simulation approach to implement the control on the population of synchronized neurons makes the control performance more accurate, minimizes the control energy consumption while achieving the desired control objective. One of the most important ways to achieve these efforts as well as the desired goals is to design a software toolbox. Various powerful software toolboxes have been designed to simulate the dynamic behavior of a neuron and networks, such as Neuron [11] and Brian [12], XPPAUT [13], and bdtoolbox [14], but to the best of our knowledge, no toolbox has ever been developed to simulate the dynamic behavior of synchronous or asynchronous neural networks, as well as to examine professional controllers to change the synchronization behavior of these networks of neurons looking through the lens of their phase distribution.

This toolbox is designed in order to investigate the theories of nouron dynamics and synchronization of (stochastic) neural populations without any special knowledge of programming and scientific simulations. The main contributions of the work are: (1) controlling the neural oscillators synchronization by phase distribution controls can be simulated without any programming efforts. (2) The proportional controlers, such as bang-bang or user-defined control inputs can be implemented. (3) There are various phase response curves (PRC) related to different neural models such as Hodgkin-Huxley (HH), Fitzhugh-Nagumo (FHN), Rose-Hindmarsh (RH) and Thalamic. (4) The dynamics of the aforementioned models can be investigated in this toolbox. (5) User-defined distributions or well-known distributions such as Von-Mises or uniform ones can be used for initial and final neurons phase distributions. (6) Different numerical approaches are developed for simulations. (7) The dynamics of neuronal populations can be determenistic or stochastic with a Gaussian white noise.

## 2. Problems and Background

The main purpose of this toolbox is to control the synchronization in populations of identical and uncoupled neural oscillators with/without noise by the phase-based control system which is introduced in [4, 15, 10]. As we mentioned earlier, this model has some remarkable advantages. Not only does it make the analysis of high-dimensional neural dynamical systems more convenient, but it also makes the designing of the control systems experimentally more applicable.

Using phase reduction and expressing the population dynamics with the probability of their distribution, the problem converts to a partial differential equation (PDE). Actually, this PDE depends on the presence or absence of noise. If there is no noise in the system, for each oscillator of the system we have:

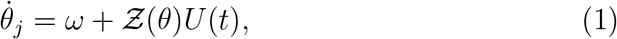

where 𝒵(*θ*) is the phase response curve (PRC) depending on the neural dynamic model [16] and *U* (*t*) is the control input. In addition, *j* = 1, …, *M* is the number of oscillators of the system. Moreover, the dynamics of the probability distribution *ρ*(*θ, t*) implied in the advection equation is as follows [4, 10]:

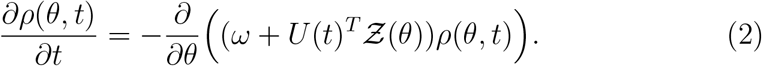

On the other hand, these equations for a noisy system are defined as:

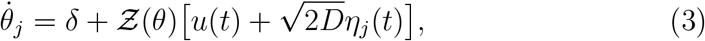

where 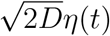 is a Gaussian white noise with zero mean and variance 2*D* affecting the control input. In addition, we consider the following equation for representing the dynamics of the probability distribution of oscillators [17, 18].

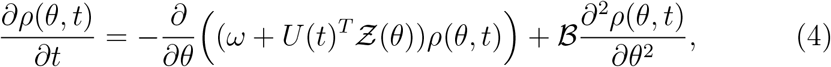

where

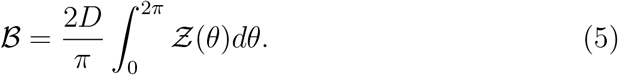

## 3. Software Framework and implementation details

The toolbox has a GUI which is designed by GUIDE in MATLAB and through which users can evaluate the synchronization or asynchrony of different neuronal populations without any programming knowledge and examine them using changing their phase distribution. This toolbox contains eighty MATLAB files and functions to provide numerous options for users in simulations including different options in numerical methods, default control algorithms, population distributions, and neural models. Moreover, there are some options that allow users to explore their models with simple user-defined MATLAB codes. NPDS toolbox includes two main parts described below: The most important and main part is neural population (De)synchronization where one can investigate distribution control models on various neural models such as HH, FHN, RH, and Thalamic neuron models. In order to evaluate the obtained results, there exist different kinds of numerical approaches for simulation. The current version (1.0) supports the 5-point stencil finite difference (FD) method, generalized Lagrange Jacobi pseudo-spectral method, radial basis function (RBF) method, radial basis function generated finite difference method (RBF-FD), and Fourier decomposition method. Moreover, we develop fourth-order Runge-Kutta algorithms to solve ODEs (1) and stochastic ODEs (3). In addition to synchronization and desynchronization, this toolbox can be used for population phase shifting and clustering which has many uses. For instance, clustering has application in rewiring neural plasticity synaptic connections between neurons. For this purpose, one can define arbitrary initial and final distributions and choose a numerical algorithm to investigate the control algorithm response. This toolbox is developed in such a way that users can follow the control strategy performance in simulation and evaluate their control algorithm in terms of accuracy, convergence, and efficiency according to the selected numerical method.

Another part of NPDS toolbox is neural dynamics which interested users in dynamical systems might be attracted to. There are some powerful computational toolkits for simulation of neural dynamical models such as Xppaut [13], Virtual Brain [19] and Brain Dynamics Toolbox [14]. NPDS toolbox users can examine the dynamics of the models introduced in the Neural population (De)synchronization part as neuronal models. They can change the parameters or initial condition of each model and see their impact on the dynamics of the problem. Moreover, there are some useful options such as showing phase portraits, adding the vector field and stream to the figures or reporting the CPU time and dynamical system state. In fact, this part complements the previous one and provides users with more complete information about the neural toolbox model. This toolbox uses standard ODE solvers (`ode23tb`, `ode45` and `ode15s`) to solve the neurons dynamical systems.

We provide a diagram (please see the figure here: https://github.com/cmplab/npds-toolbox/blob/main/docs/Pictures/Arch.png) to display the architecture of the toolbox functions and the relations between them. In this figure, the sources of graphical user interface files are represented by rectangles. We have two main part i.e `NPDSToolbox.m` and `NeuronDynamic.m`. These files are the main parts of the toolbox, which are represented by two diamonds and can be run from the command line directly. Moreover, `About.m` can be run from the command line, but it is not one of the main files of the toolbox and just gives a brief overview of the toolbox. This file is shown by a diamond in the figure. There are some main functions. These functions call the other functions to do their task correctly. On the other hand, regular functions are called by the main ones and these functions do not need to call other functions. These two types of functions are displayed by two ellipses and one ellipse, respectively. Some functions invoke simple functions defined inside the same file. Simple functions are shown by the ellipse dotted line. A diamond inside a rectangle expresses a static file that creates a user-defined function file when the user intends to define a new function. Finally, the cloud-like shape is `PARAMETER GUIDE.md` file which is a guide for model parameters.

## 4. Illustrative Examples

In this section, examples of the two main parts of the toolbox are provided. These examples are based on one of the neuronal models named the Rose-Hindmarsh (RH) model. First, we use the NeuronDynamic GUI to investigate RH neuron dynamic and show the effects of values of parameters on the state (resting, bursting, damping, and transition states) of the dynamical system. Then, in the next section, a control strategy is designed to desynchronize an RH neural population using the neural population (de)synchronization part, and we describe how to define user-defined initial distribution and control strategy.

### 4.1. Neurondynamic

The RH dynamical system is defined as follows [17, 18]

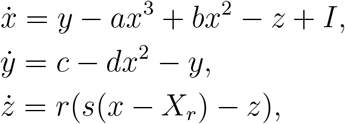

where *x* is a dimensionless variable related to the membrane potential. Also, *y* is called the spiking variable and measures the rate of transportation of sodium and potassium ions, and *z* corresponds to adaptation current. *a, b, c, d, r, s, X*_*r*_ are model parameters. Moreover, *I* is the applied current to the neuron. The model parameters in the toolbox are explicitly easily modifiable. Additionally, the initial values and the applied current can be selected from specified intervals whose bounds can be changed by the users. Note that in the toolbox toolbar, the button 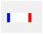 makes to change the intervals of the initial values and the applied current to the desired ones. The left and right plots in the display panel show the dynamic and phase portraits of the model, respectively. Figure 1 shows four different states including *resting, burst, damping* and *transition* by changing the model parameters and intervals. The buttons 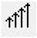 and 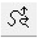 add vector field and stream to the phase portrait, used in figures 1a, 1b and 1c. Also, in Figures 1b and 1c, both portraits have grids in different sizes using buttons 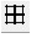 and 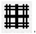. By selecting the checkboxes below the dynamic portrait, one can select the variables to show in the figure. In order to display the results more appropriately, the checkbox *scale mode* scales the behavior of the variables in the interval [0, 1] (See Figure 1c). In addition, the checkboxes below the phase portrait specify the coordinate axes of the phase portrait. At the same time, by selecting the 3*D* checkbox, the three-dimensional phase portrait of models with more than two variables can be demonstrated (see figure 1d). Finally, *neuron type, dynamic state*, and CPU time are reported in the below text box.

**Figure 1:**
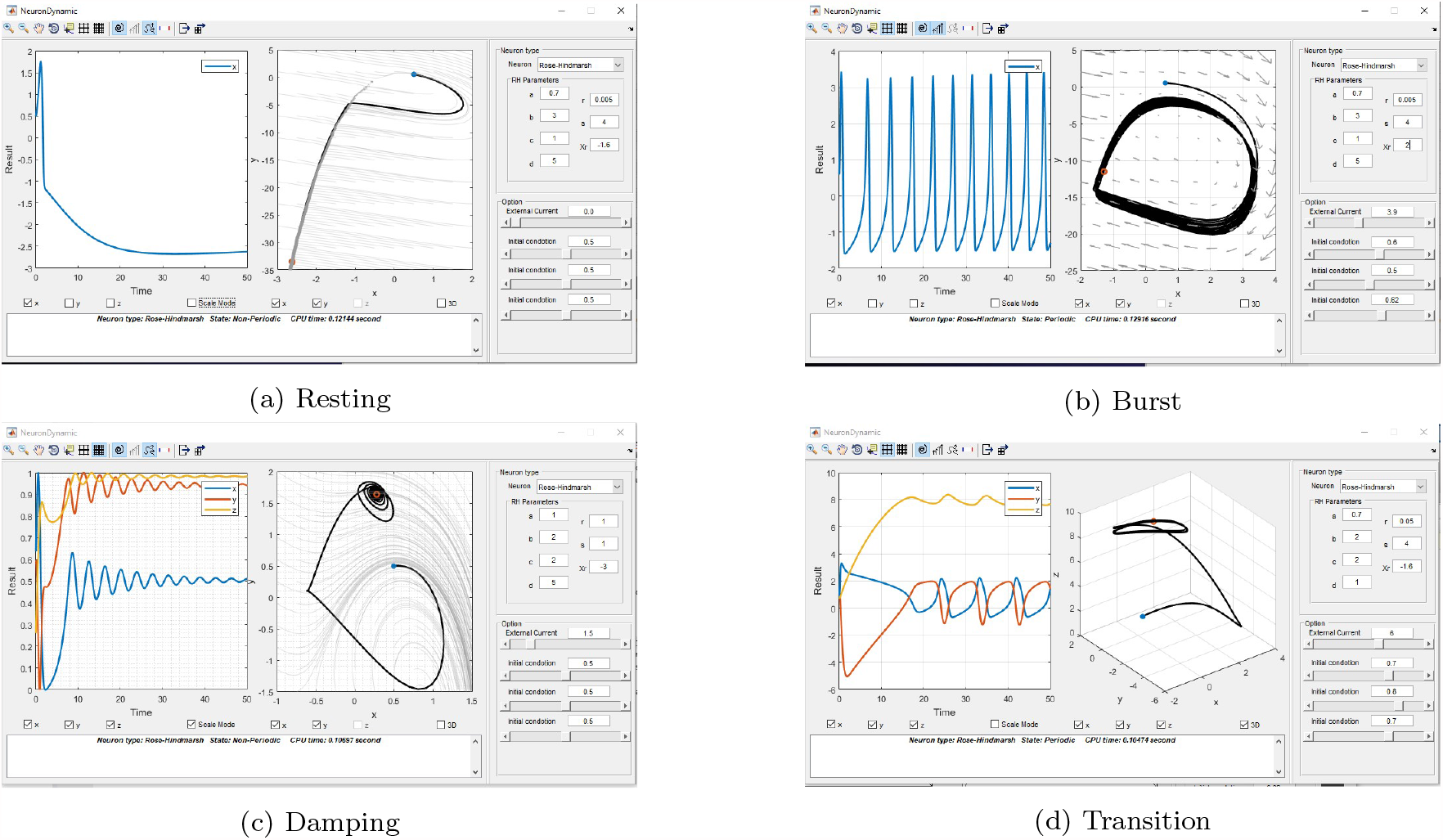
Different dynamic states of the Rose-Hindmarsh model.

### 4.2. Neural population (De)synchronization

As the next test case, we tend to desynchronize a semi-synchronous population of RH neural oscillators by a noisy bang-bang control. A semi-synchronous population of neural oscillators can be interpreted as a partial synchronization of neural oscillators in two distinct zones that are poorly connected. To apply the desynchronization process, the initial semi-synchronous population should be presented as a distribution. How to define a new distribution in the toolbox is described in the following section.

### 4.3. Defining initial and final distribution

In the toolbox, Von-Mises (single-peak) and uniform distributions are provided as defaults. However, the initial distribution in this example is equivalent to a two-peaks distribution. According to the Von-Mises distribution, the multi-peaks distributions can be like a moving wave with a constant velocity which is derived from Von-Mises distribution modification as follows [17, 18]:

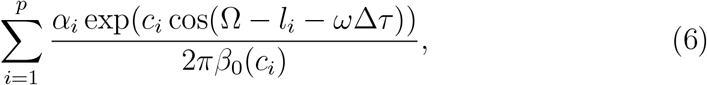

where *α*_*i*_ are average coefficients, *p* is the number of peaks, *c*_*i*_ and *l*_*i*_ are the concentration and location of each peak, *β*_0_ is the first kind of Bessel function. Note that the following condition should always behold in the distribution design

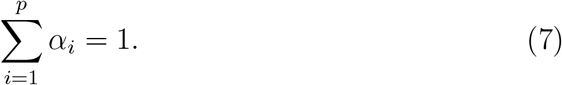

The parameters Ω, *ω*, Δ*τ, β*_0_(.) are equivalent to the keywords `doamin`, `omega`, `i*dt`, `besseli(0,.)`, respectively. In addition, in order to perform the desynchronization process, apart from the initial distribution, its derivative should also be implemented. So, the desired initial distribution and its derivative can defined as variables `dist` and `dif dist`. Consider the following example.

~~~
function [dist, diff dist]= user defined initial dist (
     domain, location, concentration, omega, i, dt)
conc =4;
alpha = 0. 5 ;
loc=pi / 10 ;
%Defining initial distribution
dist=alpha ∗exp (conc ∗ cos (domain−loc −omega∗ i ∗ dt)) /(2 ∗ pi ∗
     besseli (0, conc)) …
       +alpha ∗exp (conc ∗ cos (domain−6∗loc −omega∗ i ∗ dt)) /(2 ∗ pi
           ∗ besseli (0, conc)) ;
%Defining derivative of initial distribution
diff dist=alpha∗(−conc ∗ s i n (domain−loc −omega∗ i ∗ dt)). ∗ exp
     (conc ∗ cos (domain−loc −omega∗ i ∗ dt)) …
      /(2 ∗ pi ∗ besseli (0, conc))+alpha∗(−conc ∗ s i n (domain−6∗
        loc −omega∗ i ∗ dt)). ∗ …
exp (conc ∗ cos (domain−6∗loc −omega∗ i ∗ dt)) /(2 ∗ pi ∗
    besseli (0, conc)) ;
end
~~~

### 4.4. A user-defined control strategy

In the current version of the toolbox, the appropriate and simple control and explosion inputs can be used in a predefined way. However, in this section, we want to show how the user defines a custom controller, for example, a noisy bang-bang control input as follow

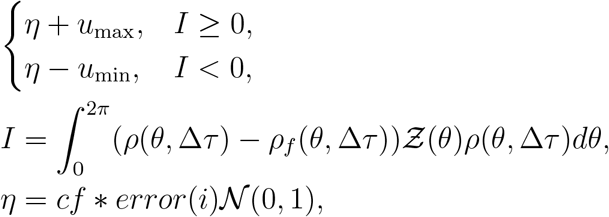

where *u*_max_ and *u*_min_ are upper and lower limits of control input, *error*(*i*) is error of result in step *i* and *cf* is a constant. To design this control we need a domain of the problem, current distribution, final distribution, PRC, current error matrices, and current step which are determined with keywords `domain`, `phi`, `phif`, `prc`, `error(.)`, and `iteration number`, respectively. Also, the desired control can define as a variable `u`.

Consider the following example:

~~~
function u=user defined control (vararg i n)
.
.
.
cf =50;
%Defining the noisy bang−bang control
I=(trapz (domain, (phi−phi_f). ∗prc ‘. ∗ phi)) ;
u max=12; % Upper limit
u min=−12; % Lower limit
r=cf ∗ error (iteration number) ∗rand (1, 1) ;
if I>0
  u=u_max+r ;
else
  u=u_min−r ;
end
~~~

Figure 2 shows the initial and final results of performing the introduced desynchronization process. According to this figure, we can evaluate the proposed control strategy that can desynchronize the neural oscillators by radial basis function generated finite difference method [18] with consuming about 1803 units of energy.

**Figure 2:**
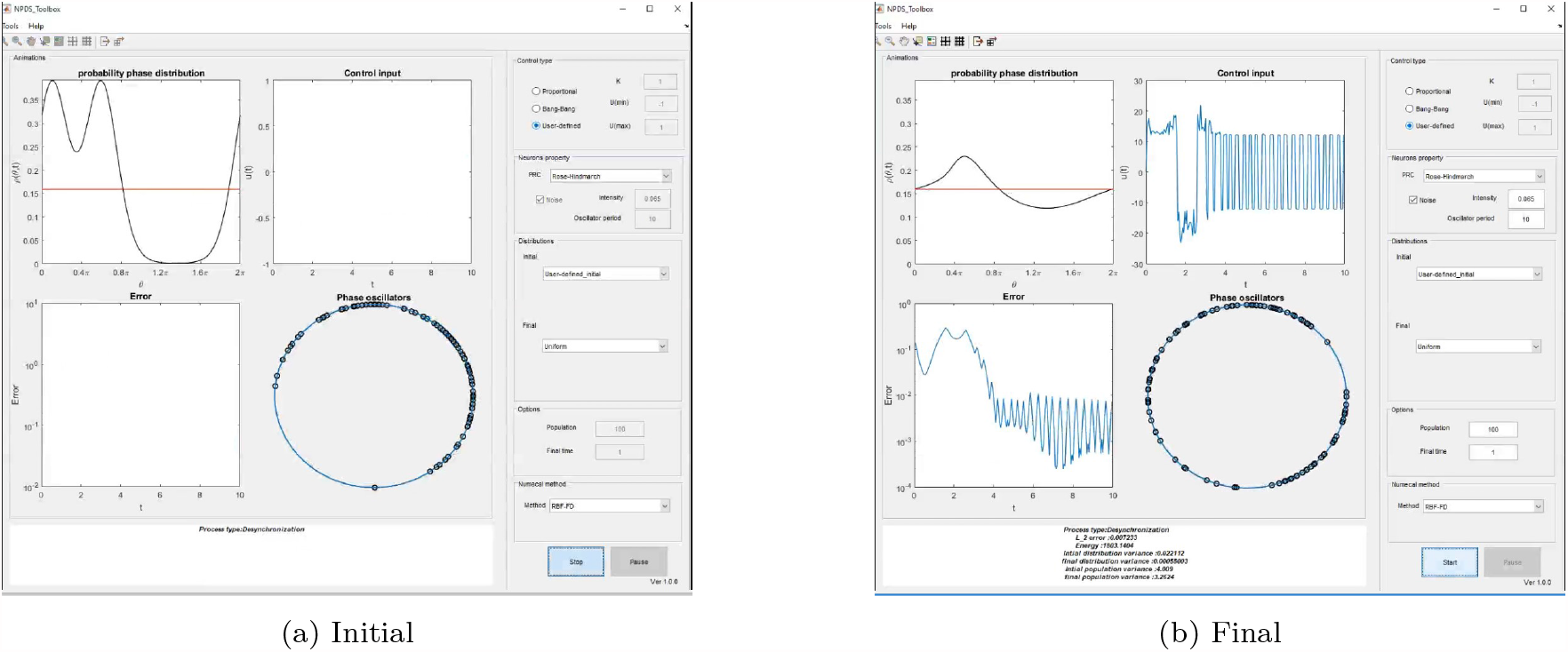
The result of applying noisy bang-bang control to desynchronize a population of neural oscillators.

## 5. Conclusions

The NPDS toolbox is an interactive graphical tool in which students/ engineers/researchers in computational neuroscience can evaluate the synchronization or asynchrony of user-defined (stochastic) neuronal populations without any programming knowledge and examine professional controllers to change the synchronization behavior of these networks of neurons looking through the lens of their phase distribution. This graphical interface imposes no limit on the size of the neural population, the type of dynamics of the neurons involved, the design of the phase change controller, nor the numerical simulation approach. The NPDS will continuously be extended with new features such as adding coupled neuronal populations as neural networks, improving numerical simulation approaches, and adding more PRCs as well as user-defined PRCs. Once any new feature is implemented, it can be easily shared with other toolbox users.

## Acknowledgements

Matlab^®^ is a registered trademark of The Mathworks, Inc., 3 Apple Hill Drive, Natick, MA 01760–2098 USA, 508-647-7000, Fax 508-647-7001, www.mathworks.com.

## Required Metadata

### Current executable software version

**Table 1:** Software metadata (optional)

| Nr. | (executable) Software metadata description |  |
| --- | --- | --- |
| S1 | Current software version | V 1.0 |
| S2 | Permanent link to executables of this version | https://github.com/cmplab/npds-toolbox |
| S3 | Legal Software License | BSD-3-Clause License |
| S4 | Computing platform/Operating System | Matlab 2016a or newer |
| S5 | Installation requirements & dependencies | None |
| S6 | If available, link to user manual - if formally published include a reference to the publication in the reference list | https://npds.readthedocs.io/en/latest/ |
| S7 | Support email for questions |  |

### Current code version

**Table 2:** Code metadata (mandatory)

| Nr. | Code metadata description |  |
| --- | --- | --- |
| C1 | Current code version | V 1.0 |
| C2 | Permanent link to code/repository used of this code version | https://github.com/cmplab/npds-toolbox |
| C3 | Legal Code License | BSD-3-Clause License |
| C4 | Code versioning system used | Github |
| C5 | Software code languages, tools, and services used | Matlab 2016a or newer |
| C6 | Compilation requirements, operating environments & dependencies | None |
| C7 | If available Link to developer documentation/manual | https://npds.readthedocs.io/en/latest/ |
| C8 | Support email for questions |  |

